# AMPK promotes Arf6 activation in a kinase-independent manner upon energy deprivation

**DOI:** 10.1101/2021.11.18.469188

**Authors:** Kuan-Jung Chen, Jia-Wei Hsu, Fang-Jen S. Lee

## Abstract

AMP-activated protein kinase (AMPK) is a crucial cellular nutrient and energy sensor that maintains energy homeostasis. AMPK also governs cancer cell invasion and migration by regulating gene expression and activating multiple cellular signaling pathways. ADP-ribosylation factor 6 (Arf6) can be activated via nucleotide exchange by guanine nucleotide exchange factors (GEFs), and its activation also regulates tumor invasion and migration. By studying GEF-mediated Arf6 activation, we elucidated that AMPK functions as a noncanonical GEF for Arf6 in a kinase-independent manner. Moreover, by examining the physiological role of the AMPK-Arf6 axis, we determined that AMPK activates Arf6 upon glucose starvation and 5-aminoimidazole-4-carboxamide-1-β-D-ribofuranoside (AICAR) treatment. We further identified the binding motif in the C-terminal regulatory domain of AMPK that is responsible for promoting Arf6 activation and thus inducing cell migration and invasion. These findings reveal a noncanonical role of AMPK in which its C-terminal regulatory domain serves as a GEF for Arf6 during energy deprivation.

## Introduction

Cells couple nutrient availability to signaling to support successful cell growth. During the decrease in energy status upon starvation, AMP-activated protein kinase (AMPK), the major energy sensor and metabolic switch, is activated to promote ATP production by upregulating catabolic pathways [1]. AMPK, a heterotrimer complex, consists of a catalytic α subunit and two regulatory β and γ subunits. The α subunit AMPK catalytic core contains an N-terminal Ser/Thr kinase domain that is activated when a conserved Thr residue at position 172 (T172) in the activation loop is phosphorylated by its upstream kinase [2]. The C-terminal regulatory domain of AMPK α binds to β. The γ subunit and appears to possess autoinhibitory properties that repress its kinase activity [3]. Emerging studies have shown that AMPK plays a regulatory role in various tumors, acting as a tumor suppressor to reduce the aggressive behaviors of cancer cells [4, 5]. Impaired regulation of AMPK activity is thought to contribute to the induction of cancer cell invasion and migration [6, 7]. In contrast, AMPK activation also contributes to cancer progression through various cellular signaling pathways [8-10]. However, the mechanism underlying AMPK-associated malignant phenotypes needs to be elucidated.

ADP-ribosylation factor 6 (Arf6), a small GTPase highly conserved across species, regulates vesicular trafficking and cytoskeletal reorganization at the plasma membrane (PM) [11, 12]. Arf6 activity is also involved in facilitating cancer cell migration and invasion [13-15]. Prevention of Arf6 cycling inhibits cancer cell progression. Mechanistically, active Arf6 is thought to act through the recruitment of distinct effectors, such as coat proteins and lipid-modifying enzymes, and through the modulation of actin structure to induce downstream cellular events [16]. Activation of Arf6 is tightly regulated by guanine nucleotide exchange factors (GEFs), which facilitate the dissociation of guanosine diphosphate (GDP) and its replacement with guanosine triphosphate (GTP). The human genome encodes 15 ARF GEFs, which are divided into six families based on sequence relatedness, domain organization, and phylogenetic analysis [17]. Several GEFs, such as ARF nucleotide-binding site opener (ARNO) containing the Sec7 domain with nucleotide exchange activity, have been identified to activate Arf6 [18, 19]. Although the Sec7 domain seems critical for promoting the activation of Arf6, we previously found that yeast Snf1/AMPK acts as a noncanonical GEF to activate yeast Arf3, a homolog of mammalian Arf6, in response to glucose depletion [20]. However, whether mammalian AMPK can also act as a noncanonical GEF for Arf6 is unknown.

In this study, we demonstrate that AMPK, unlike its canonical role with kinase activity, acts as a GEF to activate Arf6. Treatment with 5-aminoimidazole-4-carboxamide ribofuranoside (AICAR), an analog of adenosine monophosphate (AMP) that is capable of stimulating AMPK kinase activity, also promotes the activation of Arf6. Glucose depletion activates AMPK to increase Arf6 activity and thus enhances cell invasion and migration. Our data reveal an unexpected role of the AMPK-Arf6 axis in mediating cancer cell motility under glucose starvation, showing how AMPK is involved in cancer cell progression to higher-grade malignancy.

## Results and Discussion

### AMPK interacts with inactive form of Arf6

We previously identified Snf1, the yeast ortholog of mammalian AMPK, as a noncanonical GEF for the activation of Arf3, the yeast ortholog of mammalian Arf6, through its C-terminal domain [20]. We sought to further examine whether AMPK exhibits similar enzymatic activity for Arf6 activation in mammalian cells. We first characterized the interaction between Arf6 and AMPK and found that the recombinant Arf6-T27N mutant, which is thought to be trapped in the GDP-bound state, directly interacts with a C-terminal fragment of AMPK (AMPK-C) (Fig. 1A). We also confirmed that AMPK-C directly interacts with recombinant Arf6 preloaded with GDP but not GTPγS (Fig. 1B). Furthermore, the dominant-negative Arf6-T27N mutant but not the constitutively active Arf6-Q67L mutant associates with AMPK, as assessed by a coimmunoprecipitation assay. This interaction was reproduced by reciprocal immunoprecipitation (IP) using an anti-myc antibody (Fig. 1C). Taken together, these results showed that AMPK interacts directly with Arf6 in a GDP-dependent manner through its C-terminal domain.

**Fig. 1.**
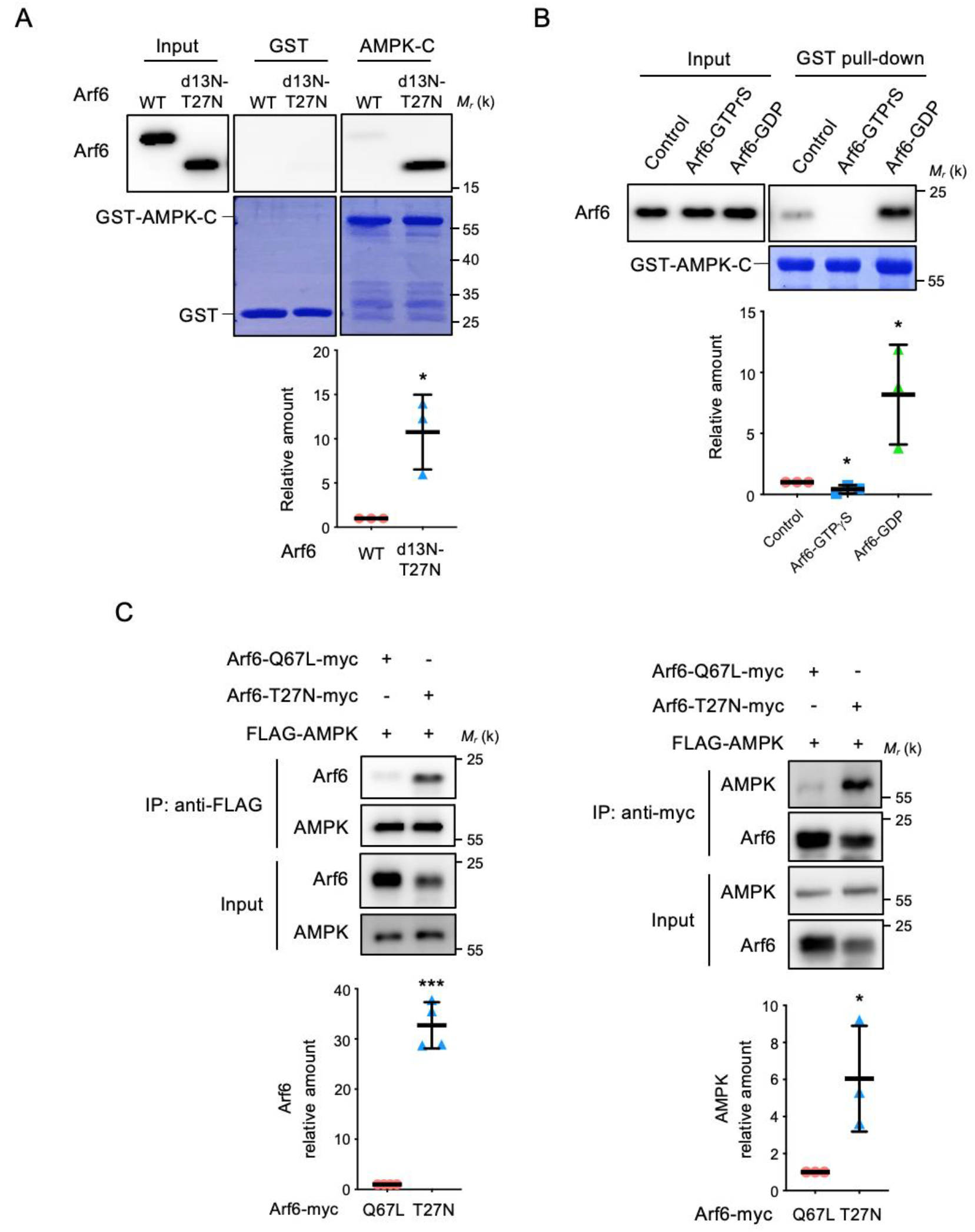
AMPK binds to the inactive form of Arf6 (Arf6-GDP) A) *In vitro* binding assay to evaluate the direct interaction between recombinant AMPK-C and Arf6. GST-AMPK-C purified from *E. coli* was incubated with recombinant His-Arf6^WT^ and His-Arf6^T27N^-d13N for 1 h. Proteins were pulled down with glutathione-Sepharose beads and visualized by immunoblotting with an anti-Arf6 antibody. The scatter plots show the mean ± s.d. values; n=3. *P≤0.05 (Student’s t-test). B) Recombinant His-Arf6 was purified from *E. coli* and loaded with GTPγS or GDP. GST-AMPK-C pulldown of Arf6 in the nucleotide-free (Control), GTP-, or GDP-bound state. Proteins were separated by SDS-PAGE and visualized by Coomassie Blue staining. The scatter plots show the mean ± s.d. values; n=3. *P≤0.05. (one-way ANOVA with Dunnett’s test). C) Coimmunoprecipitation assays confirmed the interaction between AMPK and Arf-6. 293T cells cotransfected with FLAG-AMPK and Arf6-myc plasmids (Arf6-Q67 L-myc or Arf6-T27N-myc) were lysed and subjected to IP with anti-FLAG antibodies (left panel) and anti-myc antibodies (right panel), respectively. The IP complexes were then bound to Protein A agarose beads, and the bound proteins were subjected to immunoblotting with antibodies as indicated. The scatter plots show the mean ± s.d.; n=4. *P≤0.05, ***P≤0.001 (Student’s t-test).

### C-terminal of AMPK acts as a GEF for Arf6 *in vitro*

We next sought to determine whether AMPK functions as a GEF to regulate Arf6 activation. Our previous studies showed that the C-terminal regulatory domain of Snf1 specifically activates Arf3 in yeast cells [20]. To test whether AMPK also possesses GEF activity, we examined the guanine nucleotide exchange activity of recombinant AMPK-C. We purified recombinant AMPK C-terminal fragments to examine their GEF activity toward recombinant Arf6 *in vitro* and found that AMPK-C promoted GDP release from (Fig. 2A) and GTP loading of (Fig. 2B) Arf6, comparable to the effects of cytohesin-2/ARNO (CYTH2), a well-characterized Arf6 GEF. These data indicated that AMPK-C can stimulate the release of GDP to allow the binding of GTP to Arf6.

**Fig. 2.**
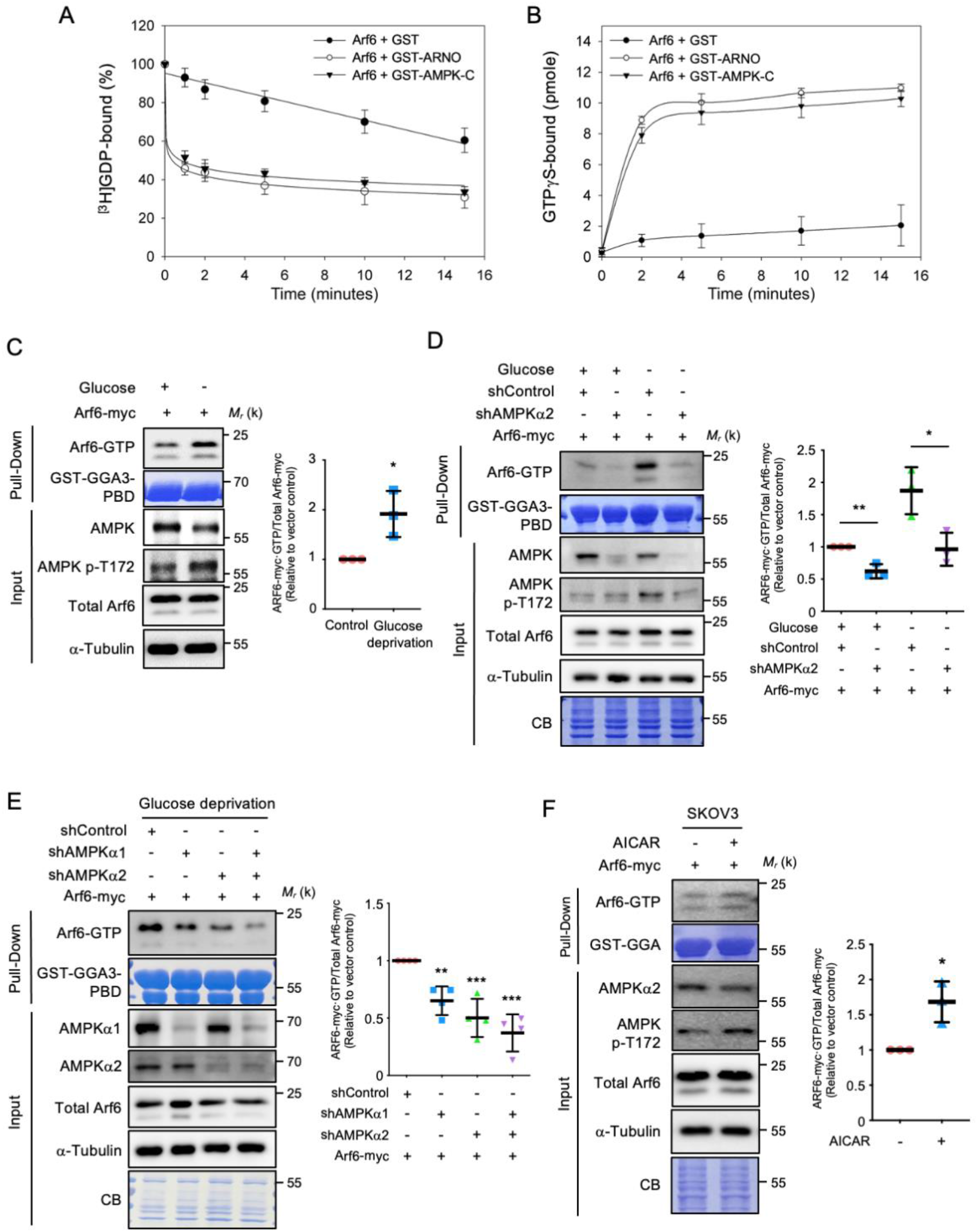
AMPK activation increases the levels of Arf6-GTP *in vitro* and *in vivo*. A, B) The C-terminus of AMPK acts as a GEF for Arf6 *in vitro*. [^3^H]GDP dissociation from (left) and [^35^S]GTPγS binding to (right) Arf6 in the presence of recombinant AMPK-C purified from *E. coli*. The data are reported as the mean ± s.d. of the percentages of dissociated [^3^H]GDP and bound [^35^S]GTPγS; n=3 (one-way ANOVA with Dunnett’s test). B) Glucose depletion increases the level of Arf6-GTP in cancer cells. SKOV3 cells were subjected to glucose deprivation for 16 h. Cell lysis followed by pulldown assays was performed to examine coexpression of the indicated plasmids. The scatter plots show the mean ± s.d. values; n=3. *P≤0.05 (Student’s t-test). C) Knockdown of AMPKα2 reduced the activation of Arf6. Pulldown assays with SKOV3 cells coexpressing the indicated plasmids or shRNAs were used to detect Arf6 activation under glucose deprivation stress. Equal amounts of GST beads were analyzed, as determined by Coomassie Blue staining. Quantification of activity by pulldown assays was based on three biological replicates. The scatter plots show the mean ± s.d. values; n=3. *P≤0.05, **P≤0.01 (one-way ANOVA with Dunnett’s test). D) Knockdown of both AMPKα1 and α2 reduced the activation of Arf6 under glucose deprivation. Pulldown assays with SKOV3 cells coexpressing the indicated plasmids or shRNAs were used to detect Arf6 activation under glucose deprivation stress. The scatter plots show the mean ± s.d. values; n=4. **P≤0.01, ***P≤0.001 (one-way ANOVA with Dunnett’s test). E) AICAR activates AMPK and increases the level of Arf6-GTP in cancer cells. SKOV3 cells were treated with AICAR for 2 h. Cell lysates were pulled down with GST-GGA3 and immunoblotted for Arf6-GTP. The scatter plots show the mean ± s.d. values; n=3. *P≤0.05 (Student’s t-test).

### AMPK activates Arf6 *in vivo*

We next considered that AMPK acts as a major coordinator of cellular energy homeostasis during glucose depletion [1]. Our previous data demonstrated that Snf1 activates Arf3 in response to glucose depletion [20]. Thus, a potential role of glucose starvation would further support the function of AMPK as a GEF for Arf6 and provide further insights into possible mechanisms by which AMPK could be regulated under this condition. Previous reports have shown that SKOV3 ovarian carcinoma cells are highly sensitive to glucose deprivation because of the induction of AMPK activity [21]. Indeed, we observed an increase in the Arf6-GTP level under glucose depletion (Fig. 2C). We investigated the involvement of AMPK in glucose starvation-regulated Arf6 activation and demonstrated a reduction in the amount of active Arf6 in cells treated with lentiviral shRNA against AMPKα2 (Fig. 2D). We also assessed the relative contributions of two AMPK catalytic isoforms, AMPKα1 and α2, to the regulation of Arf6 activation. Treating cells with lentiviral shRNA targeting either α1 or α2 decreased the amount of active Arf6, and combined inhibition of α1 and α2 resulted in a synergistic reduction in the Arf6-GTP level (Fig. 2E), suggesting that both AMPK isoforms contribute to Arf6 activation under glucose deprivation.

AMPK can be efficiently activated in cells by pharmacologic perturbation using AICAR [22]. AICAR treatment was sufficient to activate Arf6 (Fig. 2F). Phosphorylation of AMPK on T172 was also increased in cells treated with AICAR (Fig. 2F). These data further indicate that AMPK phosphorylation is correlated with Arf6 activation. Interestingly, the AMPK activator AICAR not only enhanced the kinase activity of AMPK but also stimulated its GEF activity toward Arf6. Future studies will focus on whether AICAR-induced conformational changes in AMPK assist in the exposure of the GEF domain for nucleotide exchange on Arf6.

### Activation of Arf6 by AMPK is independent of AMPK kinase activity under glucose deprivation

To further confirm the GEF activity of AMPK toward Arf6 *in vivo*, we assessed the activation status of Arf6 in cells by examining the distribution of Arf6 on the plasma membrane. Previous reports showed that Arf6 is typically distributed between the cytosolic and membrane fractions and that active Arf6 localizes predominantly to the plasma membrane [25]. Subcellular fractionation showed that glucose deprivation redistributed cytosolic Arf6 to the plasma membrane (Fig. 3A). Immunofluorescence also detected the colocalization of Arf6 and AMPK on the plasma membrane under glucose starvation (Fig. 3B). Collectively, these data indicated that AMPK acts upstream of Arf6 as a GEF to generate the GTP-bound state of Arf6. Notably, we observed that the decreased level of Arf6-GTP in AMPK-knockdown cells was rescued by the expression of wild-type (WT) AMPK (Fig. 3C). Consistent with our results in yeast [20], the expression of the kinase-dead mutant AMPK-K45R but not the phosphorylation mutant AMPK-T172A in cells activated Arf6 (Fig. 3C). These data indicated that AMPK activates Arf6 in a kinase-independent manner and that phosphorylation on Thr172 of AMPK is required for this activation process.

**Fig. 3.**
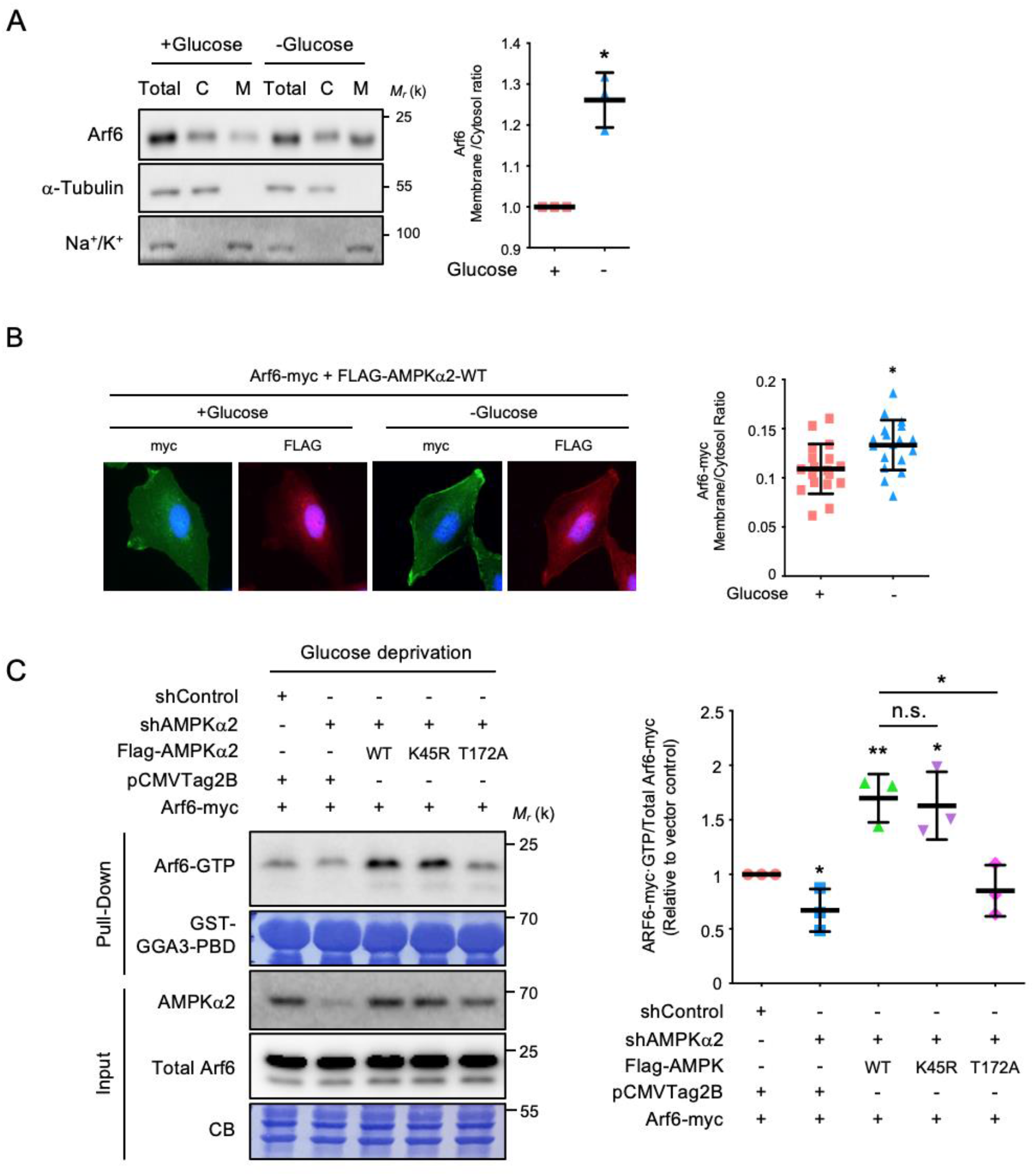
Activation of Arf6 by AMPK is independent of AMPK kinase activity under glucose deprivation. A) Glucose deprivation redistributes cytosolic Arf6 to the plasma membrane. SKOV3 cells were treated as indicated and fractionated into cytosolic (C) and membrane (M) fractions. Relative distribution of Arf6 was assessed. α-tubulin marks the cytosol, and Na^+^/K^+^-ATPase tracks the membrane. The data are presented as the mean ± s.d. values; n=3. *P≤0.05; (Student’s t-test). B) Glucose deprivation induces the colocalization of Arf6 and AMPK on the plasma membrane. SKOV3 cells were cotransfected with Arf6-myc and FLAG-AMPKα2-WT, subjected to glucose deprivation for 16 h, and then stained with anti-myc and anti-FLAG antibodies. Linear profiles, as indicated by the dashed lines created using ImageJ software, are shown for the PM and cytosolic fractions of the fluorescent proteins across the cell. The data are presented as the mean ± s.d. values; n=16. *P≤0.05; (Student’s t-test). C) Thr172 phosphorylation but not AMPK kinase activity is required for AMPK-mediated Arf6 activation under glucose deprivation. Pulldown assays were performed on 293T cells transiently transfected with Arf6-myc, ARNO, AMPK-WT, AMPK-KR and AMPK-TA. Active forms of Arf6 were pulled down with purified GST-GGA3. The GTPase was trapped in the GTP-bound state with a Golgi-associated gamma adaptin ear containing the ARF binding protein 3-glutathione-S-transferase (GGA3-GST) fusion protein coupled to Sepharose beads. Interacting proteins bound to the GST beads were analyzed by immunoblotting using the indicated antibodies. Arf6, AMPK, AMPK p-T172 and FLAG (input) in total cell lysates were analyzed by SDS-PAGE/immunoblotting to verify the initial expression levels. Equal amounts of GST beads were used in each experiment, as shown by Coomassie Blue staining. Right panel, quantitative analysis of active Arf6. The data are reported as the mean ± s.d. of three experiments compared to the empty vector (Empty) control. The scatter plots show the mean ± s.d. values; n=3. *P≤0.05; **P≤0.01; n.s., not significant. (Student’s t-test.)

### AMPK activates Arf6 through its C-terminal regulatory domain

Our previous findings demonstrated that the C-terminal α-helical hydrophobic core of Snf1 is responsible for its GEF activity toward Arf3 [20]. In addition, structural biology analysis has revealed that the C-terminal regulatory domain of AMPK is highly evolutionarily conserved, with several similar hydrophobic residues, from yeast to mammals [26]. We then generated two mutants of AMPK with substitution of those conserved hydrophobic residues with alanine: AMPK-A2 (a.a. 402-406) and AMPK-A5 (a.a. 417-421). We first found that both the AMPK-A2 and AMPK-A5 mutants were defective in binding to Arf6 (Fig. 4A). We further demonstrated that the phosphorylation of acetyl-CoA carboxylase 1 (ACC1), a known optimal substrate of AMPK, is comparable in cells expressing either AMPK-A2 or AMPK-A5 compared with cells expressing AMPK-WT (Fig. 4B), suggesting that those mutations do not alter the kinase activity of AMPK. Moreover, compared with reconstitution of AMPK-WT, reconstitution of either AMPK-A2 or AMPK-A5 in AMPK knockdown cells decreased the Arf6-GTP level (Fig. 4C). We observed that AMPK-A5 was defective in inducing Arf6 plasma membrane localization (Fig. 4D). Taken together, these findings suggested that C-terminal hydrophobic residues of AMPK interact with Arf6 to promote nucleotide exchange on Arf6. Specific sequences within the AMPK C-terminal regulatory domain contain hydrophobic residues that contribute to the Arf6 interaction, and mutations in these motifs interfered with the GEF activity of AMPK toward Arf6 but did not affect its kinase activity for substrate phosphorylation. Further structural analysis to elucidate the molecular basis of the distinct surfaces of AMPK, specifically its activation loop for kinase activity and enzymatic motif for GEF function, will be intriguing.

**Fig. 4.**
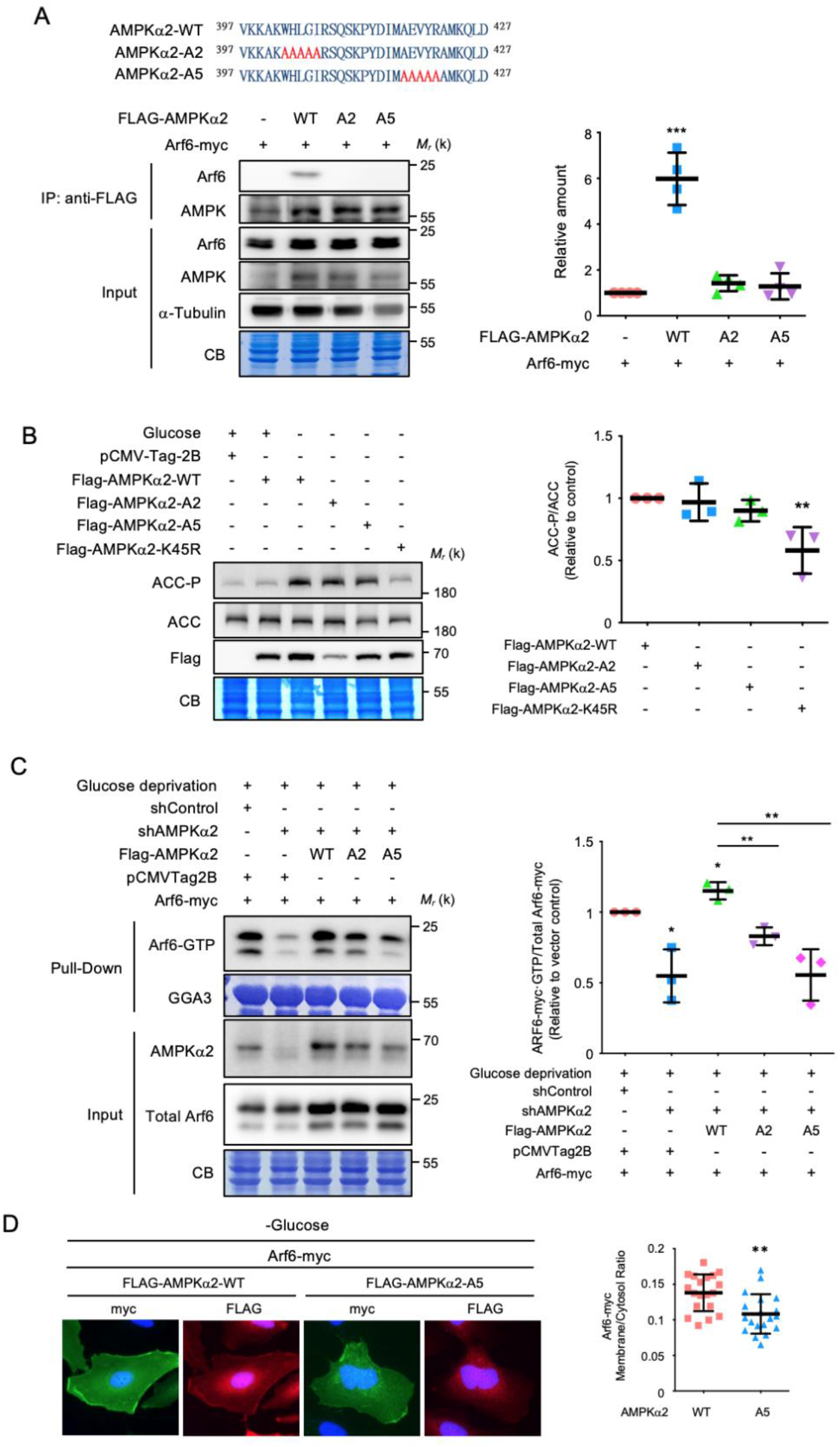
The C-terminus of AMPK interacts with and activates Arf6. A) Sequences of AMPKα2, AMPK α2-A2, and AMPK α2-A5. Interactions were verified by transiently transfecting 293T cells with the indicated plasmids, performing IP with an anti-FLAG antibody, and immunoblotting the bound proteins to detect Arf6. The scatter plots show the mean ± s.d. values; n=3. B) The A2 and A5 mutants did not affect the kinase activity of AMPK. 293T cells were transfected with the indicated plasmid for 24 h and then treated with AICAR for 16 h. Cells were harvested, and the ACC and ACC phosphorylation levels were checked by western blot analysis. The scatter plots show the mean ± s.d. values; n=3. **P≤0.01 (one-way ANOVA with Dunnett’s test). C) AMPK-A2 and AMPK-A5 failed to rescue the reduced Arf6-GTP level in AMPK-knockdown cells. Pulldown assays were performed on AMPKα2-knockdown 293T cells with expression of AMPKα2, AMPK α2-A2 and AMPKα2-A5, as indicated. Right, quantitative analysis of active Arf6. The scatter plots show the mean ± s.d. values; n=3. *P≤0.05; **P≤0.01 (one-way ANOVA with Dunnett’s test). D) A wild-type but not Arf6-binding defective mutant induces the plasma membrane localization of Arf6 under glucose deprivation. SKOV3 cells were knocked down with AMPKα2 and then cotransfected with Arf6-myc and FLAG-AMPKα2-WT or A5 mutant. Cells were subjected to glucose deprivation for 16 h and then stained with anti-myc and anti-FLAG antibodies. The data are presented as the mean ± s.d. values; n=19. **P≤0.01 (Student’s t-test).

### A critical role for AMPK-mediated activation of Arf6 in cancer cell migration

Activation and high expression of Arf6 correlate strongly with invasion and metastasis in several tumor types [27, 28]. In addition, both a tumor suppressive and oncogenic role of AMPK has been reported [8, 29]. We finally tested whether interfering with the ability of AMPK to act on Arf6 by modulating its GTP loading ability affects cancer cell invasion. For this purpose, we performed a Matrigel invasion assay in SKOV3 cells with knockdown of the identified Arf6 GEF ARNO as invasion control cells (Fig. 5A). Cancer cell migration across a Transwell membrane was then measured to assess cell motility. Knockdown of AMPK enhanced cell invasion under normal glucose conditions (Fig. 5A). Consistent with a previous study, glucose deprivation enhanced the invasion of SKOV3 cells, but cell invasion was blocked by loss of AMPK under glucose starvation [30]. These data collectively indicate that energy stress is involved in switching the role of AMPK in the modulation of cell invasion. To investigate whether this effect on invasion is mediated through activation of Arf6 by AMPK, we ectopically expressed AMPKα2-WT to rescue cell invasion in AMPK-knockdown SKOV3 cells under glucose deprivation. Importantly, expression of Arf6-binding detective mutants of AMPK, AMPK α2-A2 and AMPK α2-A5 failed to restore this reducing invasion abilities (Fig. 5B). In addition, a decrease in wound closure was observed when AMPK α2-A2 and A5 were used to rescue AMPK knockdown cells under glucose deprivation (Fig. 5C). These data indicated that activation of Arf6 by AMPK is critically important to promote cancer cell migration specifically under glucose deprivation conditions.

**Fig. 5.**
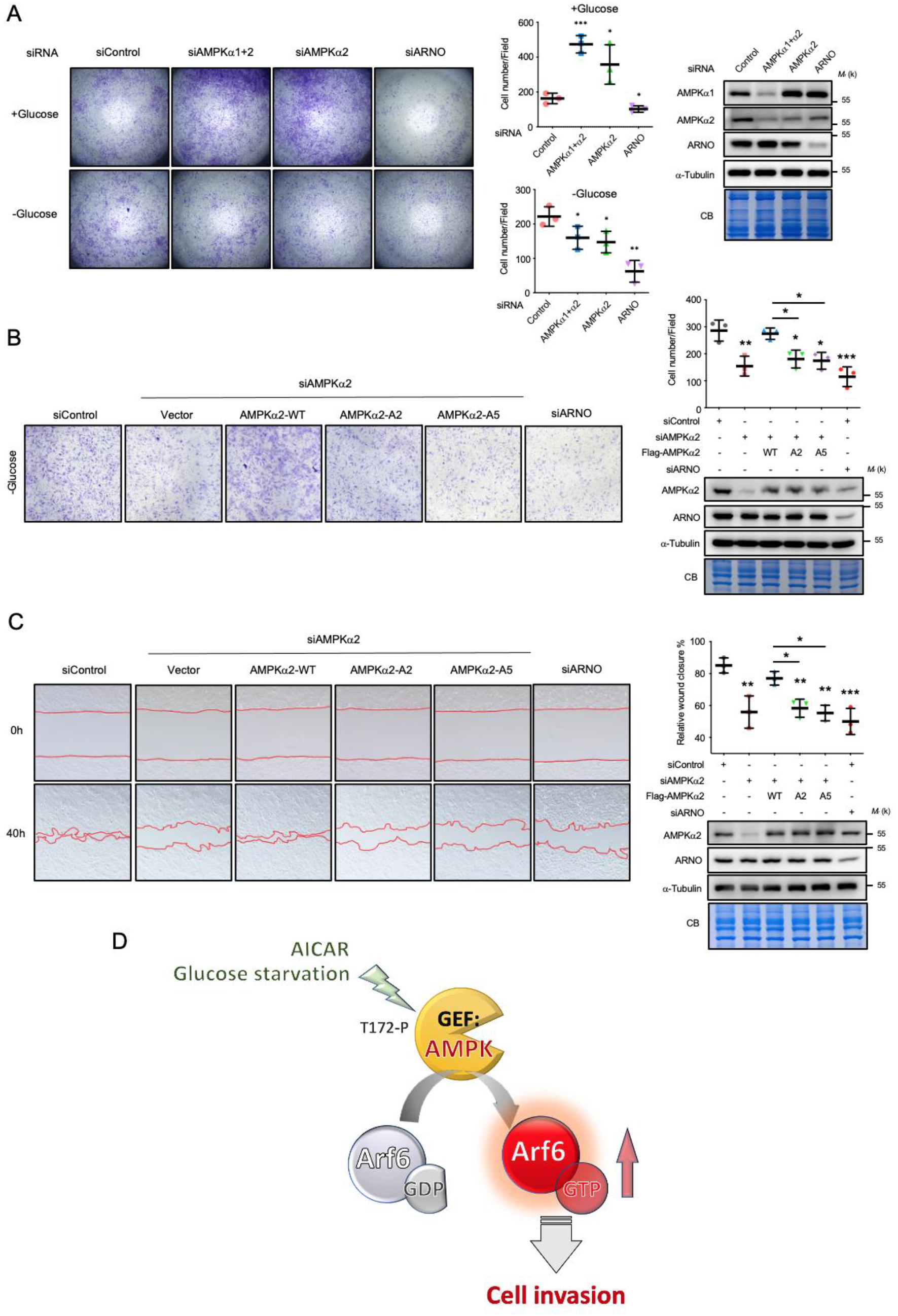
The AMPK-Arf6 axis is necessary for glucose deprivation-induced cell invasion. A) AMPK knockdown enhances the cellular invasion ability under normal glucose conditions. Transwell invasion assays were performed with SKOV3 cells after knockdown with different siRNAs for 48 h. The upper surface of the Transwell membranes was coated with Matrigel. Then, cells were seeded in the upper chambers of the Transwell cell culture apparatus. After 24 h of cell invasion, cells were stained with crystal violet for visualization and quantification. Protein expression levels were checked by immunoblotting. The graph shows the resulting quantification, presented as the mean number of cells per photographic field. The scatter plots show the mean ± s.d. values; n=3. *P≤0.05; **P≤0.01; ***P≤0.001 (one-way ANOVA with Dunnett’s test). B) AMPK-A2 and AMPK-A5 failed to rescue the reduced invasion abilities in AMPK-knockdown cells under glucose deprivation. Representative images of invasion assays of SKOV3 cells transfected with the indicated plasmids or siRNA. The scatter plots show the mean ± s.d. values; n=3. *P≤0.05; **P≤0.01; ***P≤0.001 (one-way ANOVA with Dunnett’s test). C) AMPK-A2 and A5 fail to rescue wound healing under glucose deprivation. Representative images of wound healing assays of SKOV3 cells transfected with the indicated siRNAs and plasmid under glucose deprivation. Confluent monolayers were photographed at 40 h after wounding. The scatter plots show the mean ± s.d. values; n=3. *P≤0.05; **P≤0.01; ***P≤0.001 (one-way ANOVA with Dunnett’s test). D) A model of Arf6 activation by AMPK in response to glucose deprivation. AMPK T172 phosphorylation stimulated by AICAR and glucose deprivation induces AMPK activation. AMPK directly interacts with the GDP-bound form of Arf6 and facilitates GDP dissociation and GTP loading. The AMPK-Arf6 pathway enhances the invasion ability of cancer cells.

Most previous data suggest that AMPK could be a tumor suppressor due to functional loss of an AMPK upstream kinase, liver kinase B1 (LKB1) [31]. However, recent accumulated data also indicate that AMPK might be a protumorigenic regulator. For instance, AMPK activation by lysophosphatidic acid (LPA) promotes the metastasis of SKOV3 cells [29]. Energy stress in cells due to glucose depletion appears to be a driving force of cancer cell invasiveness and migration [30]. Specifically, AMPK is usually activated when T172 is phosphorylated by an upstream kinase in response to glucose starvation. We found that the oncogenic function of AMPK may be correlated with its downstream substrate Arf6, as under these conditions, AMPK acts as a GEF rather than a kinase of Arf6 (Fig. 5D). The paradox between the protective role of AMPK in metabolic adaptation and its tumorigenic role in promoting cancer cell motility during energy imbalance needs further investigation.

## Conclusion

In this study, we revealed that AMPK functions as a GEF to directly activate Arf6, leading to the redistribution of cytosolic Arf6 to the plasma membrane, which then promotes cancer cell invasion and migration. This study shows a fundamentally different role of AMPK as a GEF rather than a kinase of its downstream substrate. Thus, AMPK, the major energy sensor, is a potential anticancer target, indicating that inhibiting the GEF activity of AMPK may decrease cancer cell motility in response to glucose starvation.

## Methods

### Plasmids and cell culture

Human cervical carcinoma HeLa cells were purchased from the American Type Culture Collection (ATCC; Manassas, VA, USA). The human serous cystadenocarcinoma cell line SKOV3 was kindly provided by Dr. Lin-Hung Wei, National Taiwan University Hospital, Taipei City, Taiwan. HEK 293T cells were provided by Dr. Tzuu-Shuh Jou, National Taiwan University, Taipei City, Taiwan. All cells were cultured in high-glucose Dulbecco’s modified Eagle’s medium (HG-DMEM; SH30007.02, HyClone) supplemented with 10% fetal bovine serum (FBS; HyClone) at 37°C in an atmosphere containing 5% CO_2_. Cells were transiently transfected with the indicated plasmids or siRNAs using Lipofectamine 2000 transfection reagent (Invitrogen) according to the manufacturer’s instructions. The shControl and shAMPK (TRCN002168) lentiviral pLKO.1 plasmids were provided by the National RNAi Core Facility of Academia Sinica, Taipei, Taiwan. The AMPKα2 K45R and T172A plasmids were gifts from Dr. Sheng-Chung Lee, National Taiwan University, Taipei, Taiwan. For expression as a GST fusion protein in *E. coli* and an N-terminal FLAG fusion protein in mammalian cells, ARNO was cloned into the pGEX-4T and pCMV-Tag2 vectors as described previously. The GGA3-PBD (1-316 a.a.) cDNA was amplified from a HeLa cell cDNA pool [32].

### *In vitro* binding assay

*E. coli* strain BL21 was transformed with plasmids, including pET32a-Arf6 (WT or d13N-T27N). After cells were induced with 0.5 mM isopropyl β-D-1-thiogalactopyranoside (IPTG) at 37°C for 3 h, glutathione S-transferase (GST) fusion proteins and His-tagged proteins were purified from

*E. coli* lysates using glutathione-Sepharose 4B (GE Healthcare Amersham, Piscataway, NJ) and nickel affinity resin (Qiagen, Valencia, CA), respectively. In the pulldown assays, GST and GST-AMPK-C (291-552 a.a.) were immobilized on glutathione beads and incubated with His-Arf6 (WT or T27N) in binding buffer containing 25 mM Tris-HCl (pH 7.5), 150 mM NaCl, 0.5% Nonidet P-40 (NP-40), 5 mM MgCl_2_, and 1× protease inhibitor for 1 h at 4°C. Then, the beads were washed three times with 1 ml of binding buffer. Next, bound proteins were analyzed by western blotting using anti-His monoclonal antibodies (BD Biosciences, La Jolla, CA, no. 552565; 1:5,000). For the GTP and GDP loading assay, WT Arf6 was loaded with either GDP or GTP by incubation with 1 mM GDP or nonhydrolyzable GTPγS, respectively, at 30°C for 15 min. The binding assay was then performed as described above.

### *In vitro* GEF activity assays

Guanine nucleotide exchange assays were performed by measuring the binding of GTPγS. In brief, 1 mM Arf6 protein and 8 mM [^35^S]GTPγS in exchange buffer containing 50 mM HEPES (pH 7.5), 1 mM MgCl_2_, 100 mM KCl and 1 mM dithiothreitol (DTT) were incubated with 100 nM GST or GST-AMPK-C. Samples were collected at various time points, diluted with 1.2 ml of ice-cold stopping buffer (50 mM HEPES (pH 7.5), 10 mM MgCl_2_, 100 mM KCl and 1 mM DTT) and filtered on membrane filter nitrocellulose filters (Millipore, AAWP04700). The filters were washed, dried and counted in a liquid scintillation counter (Beckman, LS6000IC). For the GDP dissociation assays, His-tagged ARF protein (1 mM) was first loaded with 20 mM [^3^H]GDP for 50 min at 30°C in exchange buffer. Spontaneous nucleotide dissociation and exchange were catalyzed by the addition of 1 mM unlabeled GTP and 100 nM GST, GST-AMPK-C or GST fusion proteins containing various SNF1 fragments. After incubation for various times, proteins and bound nucleotides were isolated by filtration through nitrocellulose filters and counted in a Beckman LS 5000 scintillation counter to quantify the amount of [^35^S]GTPγS and [^3^H]GDP-bound protein.

### Small G protein activity pulldown assay

For the Arf6 activity assay, SKOV3 cells were transfected with the indicated plasmid or knockdown lentiviral plasmid. For glucose deprivation treatment, SKOV3 cells were prewashed with phosphate-buffered saline (PBS) three times. Then, glucose-free DMEM (Gibco, 11966-025) containing 5% FBS was added for 16 h. Cells were lysed in pulldown buffer containing 25 mM Tris-HCl (pH 7.5), 150 mM NaCl, 0.5% NP-40, 5 mM MgCl_2_, and 1× protease inhibitor. A 40-μg sample of GST-GGA3 was incubated with 1 mg of cell lysate for 1 h at 4°C. The GST beads were then washed three times with pulldown buffer. Proteins were eluted with protein loading buffer and analyzed by immunoblotting.

### Coimmunoprecipitation

293T cells were seeded in 10 cm dishes and transiently transfected with the indicated plasmid using Lipofectamine 2000 transfection reagent. After 16 h of transfection, the cells were lysed with IP buffer (50 mM Tris-HCl (pH 7.4), 150 mM NaCl, 1% NP-40, 1× protease inhibitor cocktail), and the lysate was centrifuged at 14,000 ×g for 10 min at 4°C. The supernatants were incubated with a monoclonal anti-myc antibody (Covance, MMS-150R) or anti-Flag antibody (Sigma, F3165) for 2 h at 4°C. Then, the cells were incubated with Protein A Sepharose beads (Merck, GE17-5280-01) for 1 h at 4°C. The beads were washed three times with wash buffer. The coimmunoprecipitated proteins were eluted with sample buffer, boiled and analyzed via western blotting.

### Transwell invasion assay

The Transwell invasion assay was performed using Boyden Transwell chambers with 8-μm pore size polycarbonate membranes (Corning, costar, #3464). Transwell membranes were precoated with 70 μl of Matrigel (diluted 1:5 with culture medium, Corning, #356234) in a 24-well Transwell plate, and the Matrigel was allowed to solidify at 37°C for 1 h. After transfection of the indicated plasmids, 6×10^4^ cells in serum-free DMEM containing specific glucose concentrations were seeded into the prepared upper Transwell chambers, and 500 μl of DMEM supplemented with 10% FBS was added to the lower chambers. The 24-well plate was then incubated at 37°C in 5% CO_2_ for 24 h. Each assay was performed in duplicate. Cells were fixed with 4% paraformaldehyde/PBS for 15 min at room temperature. Noninvaded cells on the upper surface of the membranes were removed with a cotton swab, and invaded cells attached to the lower surface were stained with 1% crystal violet for 15 min. After washing, the membranes were air dried, and the invaded cells were imaged using a phase contrast microscope (Nikon, Eclipse TS-100) connected to an imaging system (Nikon, DS-5 M). The invaded cells in at least three fields per well were counted using ImageJ software.

### Total protein extraction and immunoblot analysis

Western blotting was performed as described previously [33]. After washing, the membranes were developed using an enhanced chemiluminescence system (Millipore). α-Tubulin was used as the internal control for protein loading. GST-tagged proteins were visualized by Coomassie Blue staining. Anti-α-tubulin (1:5,000, T5168; Sigma-Aldrich), anti-AMPKα1 (1:1,000, 2795S; Cell Signaling), anti-AMPKα2 (1:1,000, 2757S; Cell Signaling), anti-CYTH2/ARNO (1:1,000, H00009266; Abnova), anti-Arf6 (1:1,000, #sc-7971, Santa Cruz), anti-AMPK-P (1:1,000, 2535S; Cell Signaling), anti-myc (1:1,000, MMS-150R; Covance), and anti-FLAG M2 (1:1,000, F-3165; Sigma-Aldrich) antibodies were used.

### Immunofluorescence staining

The cells were fixed with 4% paraformaldehyde in PBS for 15 min. After washing with PBS, the cells were permeabilized with 0.01% Triton X-100 in PBS at room temperature for 5 min and blocked with 0.5% BSA in PBS. Primary antibodies were diluted in blocking buffer at the following ratios: anti-myc (1:200) and anti-FLAG M2 (1:200). Alexa Fluor 594 (A-11012; Invitrogen; anti-rabbit)- and 488 (A-11001; Invitrogen; anti-mouse)-conjugated secondary antibodies were used at a dilution of 1:1,000. Images were acquired using an Axioplan microscope (Carl Zeiss, Inc.). Quantification of the membrane-targeting PM/C ratio followed a previous method [33].

### Wound healing assay

For the wound healing cell migration assay, cells were seeded in culture medium in six-well plates at a density of 8 × 10^5^ cells per well under glucose deprivation conditions. The layer of confluent cells was scratched using a fine pipette tip, washed twice with medium, and observed using a microscope (Carl Zeiss) after 22 h. The cell migration capacity was determined using ImageJ software.

### Cell fractionation assay

SKOV3 cells (1 × 10^7^) were collected and washed twice with PBS and were then resuspended in 400 ml of buffer that consisted of 25 mM HEPES (pH 7.4), 2.5 mM MgCl_2_, 250 mM sucrose, and protease inhibitor. Cells were homogenized by passing through a 30-gauge needle 20 times. In general, 90-95% of nuclei were released from the cells, as determined by microscopy. The resulting homogenate was centrifuged at 100× g for 10 min to obtain the postnuclear supernatant. This supernatant was further centrifuged at 22,000 × g for 1 h (rotor) to obtain the cytosolic (supernatant) and membrane (pellet) fractions. Equal amounts of both fractions were analyzed by western blot.

### Lentiviral knockdown

AMPKα1 and α2 knockdown by lentiviral shRNA silencing was performed by using packaging plasmids to produce lentiviruses containing pLKO.1-GFP-shRNA vectors (RNAi Core, Academia Sinica, Taiwan). After 24 h of infection, cells were selected with puromycin for 3 days. Green fluorescent protein (GFP) was used as a reporter for viral infection, and GFP-positive cells were evaluated to determine the infection efficiency. The sequences of the siRNAs and shRNAs used in this study are listed in Table S1.

### Statistical analysis

All data are expressed as the mean ± s.d. values, and P-values were calculated by two-tailed Student’s t-test or one-way analysis of variance (ANOVA) followed by Dunnett’s post hoc multiple comparison test using Excel or Prism 8. Significant differences (*P≤0.05; **P≤0.01; ***P≤0.001) are indicated. Each independent *in vitro* experiment included three biological replicates.

## Acknowledgments

We thank Drs. Joel Moss and Randy Haun for their critical review of this manuscript. We also thank the staff of the biomedical resource center at the First Core Labs, National Taiwan University College of Medicine for technical assistance.

## Competing Interests

The authors have no competing financial interests to declare.

## Author Contributions

K.-J C., J.-W. H. and F.S.L. designed the study and interpreted the results; K.-J C. and J.-W. H. performed the majority of the experiments and analyzed the data; K.-J C. prepared the draft of the manuscript; K.-J C., J.-W. H. and F.S.L. wrote and edited the manuscript; and F.S.L. provided supervision, acquired funding, and performed project administration.

## Funding

This work was also supported by grants from the International Cooperation Project National Taiwan University and the Center of Precision Medicine from the Featured Areas Research Center Program (2020) within the framework of the Higher Education Sprout Project of the Ministry of Education in Taiwan awarded to F.S.L. This work was also supported by grants from the Ministry of Science and Technology, Taiwan (109-2636-B-002-015- and 110-2636-B-002-012-to J.-W. H.), and by the Yushan Young Scholar Program from the Ministry of Education to J.-W. H.

---

**Table S1.**
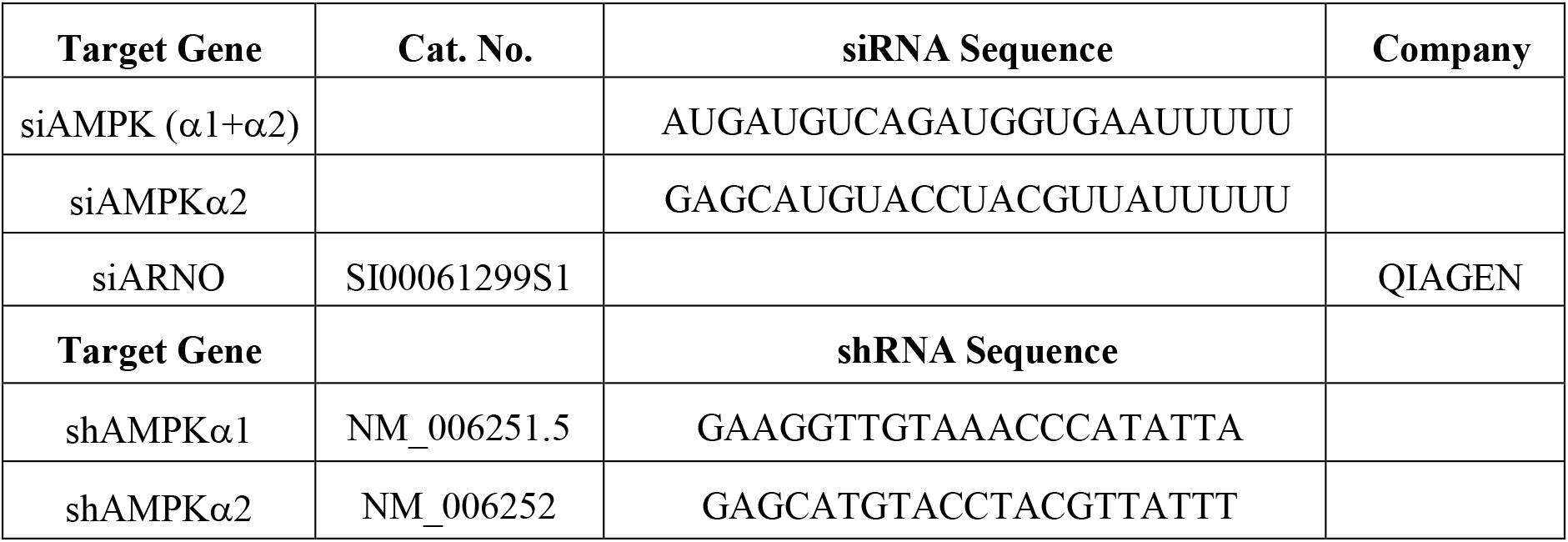
Information of siRNA and shRNA oligonucleotides

## Notes

### Competing Interest Statement

The authors have declared no competing interest.

